# First record of a Nathusius’ pipistrelle (*Pipistrellus nathusii*) overwintering at a latitude above 60°N

**DOI:** 10.1101/2020.02.24.962563

**Authors:** AS Blomberg, V Vasko, S Salonen, G Pētersons, TM Lilley

**Affiliations:** Department of Biology, University of Turku, Turku, Finland; Finnish Museum of Natural History, University of Helsinki, Helsinki, Finland; Latvian University of Life Sciences and Technologies, Jelgava, Latvia

**Keywords:** Migration, Hibernation, Range expansion, Chiroptera, Climate change

## Abstract

Highly mobile species are considered to be the first to respond to climate change by transforming their ranges of distribution. There is evidence suggesting that *Pipistrellus nathusii*, a long-distance migrant, is expanding both its reproduction and overwintering ranges to the North. We recorded the echolocation calls of bats at 16 sites in South-Western Finland on two consecutive winters, and detected calls of *P. nathusii* at one of the sites throughout the latter winter. To our knowledge, this is the northernmost observation of an overwintering *P. nathusii*, and further evidence that the species is already responding to climate change.

---

Climate change is already affecting the range limits and phenology of organisms across a plethora of taxa (Parmesan and Yohe 2003; Hickling et al. 2006; Tingley et al. 2009). According to a review by Thomas (2010), it is possible that more than half of observed animal range boundaries have shown a response to climate change. In most cases, range expansion has occurred on the edge of their of their distribution range with lower ambient temperatures, which appears to take place more rapidly than local extinctions at the side with higher ambient temperatures (Hickling et al. 2006; Thomas et al. 2006; Brommer et al. 2012). While it is difficult to differentiate changes caused by climate change from those resulting from habitat loss or other local changes, or an increase in sampling effort, there is some evidence that bats are also already responding to the changing climate through range expansion (Lundy et al. 2010; Sherwin et al. 2013). In Europe, *Pipistrellus kuhlii* (Sachanowicz et al. 2006; Ancillotto et al. 2016) and *Hypsugo savii* (Lehotská and Lehotský 2006; Uhrin et al. 2016) have shown a remarkable increase in their geographical range, whereas in North-America, Willis and Brigham (2003) reported the tree roosting *Lasiurus borealis* in Southwestern Saskatchewan, Canada, 300 km from the nearest previous observation.

*Pipistrellus nathusii* is a long-distance migrant, with distances of up to 1905 km between its breeding and overwintering areas (Pētersons 2004). Highly mobile species, such as *P. nathusii*, are most likely to be the first to respond to a changing climate by shifting their ranges towards the poles or to higher latitudes (Warren et al. 2001; Brommer et al. 2012). Indeed, Lundy et al. (2010) showed that *P. nathusii* has already adapted its distribution range in Europe. Breeding colonies, and a likely increase in the amount of individuals, have been documented in the British Isles (Russ et al. 2001; Matthews et al. 2018), Northern Italy (Martinoli et al. 2000) and on the Iberian Peninsula (Flaquer et al. 2005).

Winter is often a critical time for animals inhabiting high latitudes. Therefore, for many relatively sedentary species, such as most hibernating mammals, climate change can have unfavorable effects (Lane et al. 2012). Bats, however, are mobile and more capable of shifting their overwintering ranges as a response. Indeed, Humphries et al. (2002) suggest that the hibernation ranges of North American bats will shift towards the north because of climate change. Furthermore, according to Newson et al. (2009), climate change is likely to affect the hibernation site selection and species composition in hibernacula to an extent that the abundance of bats, and changes in the distribution and species composition at underground hibernation sites can be used as an indicator for climate change. Haarsma et al. (2019) discovered that *Myotis dasycneme* males have altered their overwintering areas, suggesting that hibernating closer to breeding areas extend their mating season while also saving energy needed for migration.

*P. nathusii* usually hibernates in solitary or in small groups above ground (Sachanowicz et al. 2019). Typical winter roosts are buildings, hollow trees, woodpiles and sometimes rock crevices (Diez and Kiefer 2016). Therefore, they are seldom observed in hibernation surveys and most findings in the winter occur by accident (Sachanowicz et al. 2019). The species overwinters predominantly in Western, Central and Southern Europe. However, there is an increasing number of observations of hibernating individuals in Eastern Europe, most notably from Poland, Slovenia, Slovakia and Hungary (Benda and Hotový 2004; Sachanowicz and Ciechanowski 2006; Sachanowicz et al. 2019; Nusová et al. 2019). Furthermore, one individual was discovered hibernating in a crevice between the window frame and concrete wall in Riga, Latvia, on 16.01.2014. Sachanowich et al. (2019) stated that *P. nathusii* might be utilizing urban heat islands to extend their overwintering range towards North-Eastern Europe. As a result, the breeding and overwintering areas of *P. nathusii* are now overlapping in Central-Europe (Sachanowicz et al. 2019).

While climate change may lead to loss in biodiversity especially in Southern Europe, many studies predict that species richness in the Northern Europe and Scandinavia will increase (Levinsky et al. 2007; Rebelo et al. 2010). According to Mikkonen et al. (2015), the mean annual temperatures in Finland have risen 2.3 ± 0.4 °C from the mid-19th century. The highest increases in temperature have taken place over the winter months, most notably in December when the monthly mean temperature has risen 4.8 °C. Furthermore, the spring months have also warmed more than the annual average. Until recently, *P. nathusii* has been considered a vagrant in Finland during migration periods in late May and late August-early October (Rydell et al. 2014; Ijäs et al. 2017). The first breeding colony was found in Southern Finland in 2006 (Hagner-Wahlsten and Kyheröinen, 2008). Since then, observations of *P. nathusii* have become more frequent outside migration period and more breeding colonies have been discovered (Hagner-Wahlsten and Karlsson 2009; Eeva-Maria Tidenberg, personal communication; Blomberg, Vasko and Lilley, unpublished data). However, until now, the species has never been recorded in the area during the hibernation period (Figure 1).

**Figure 1.**
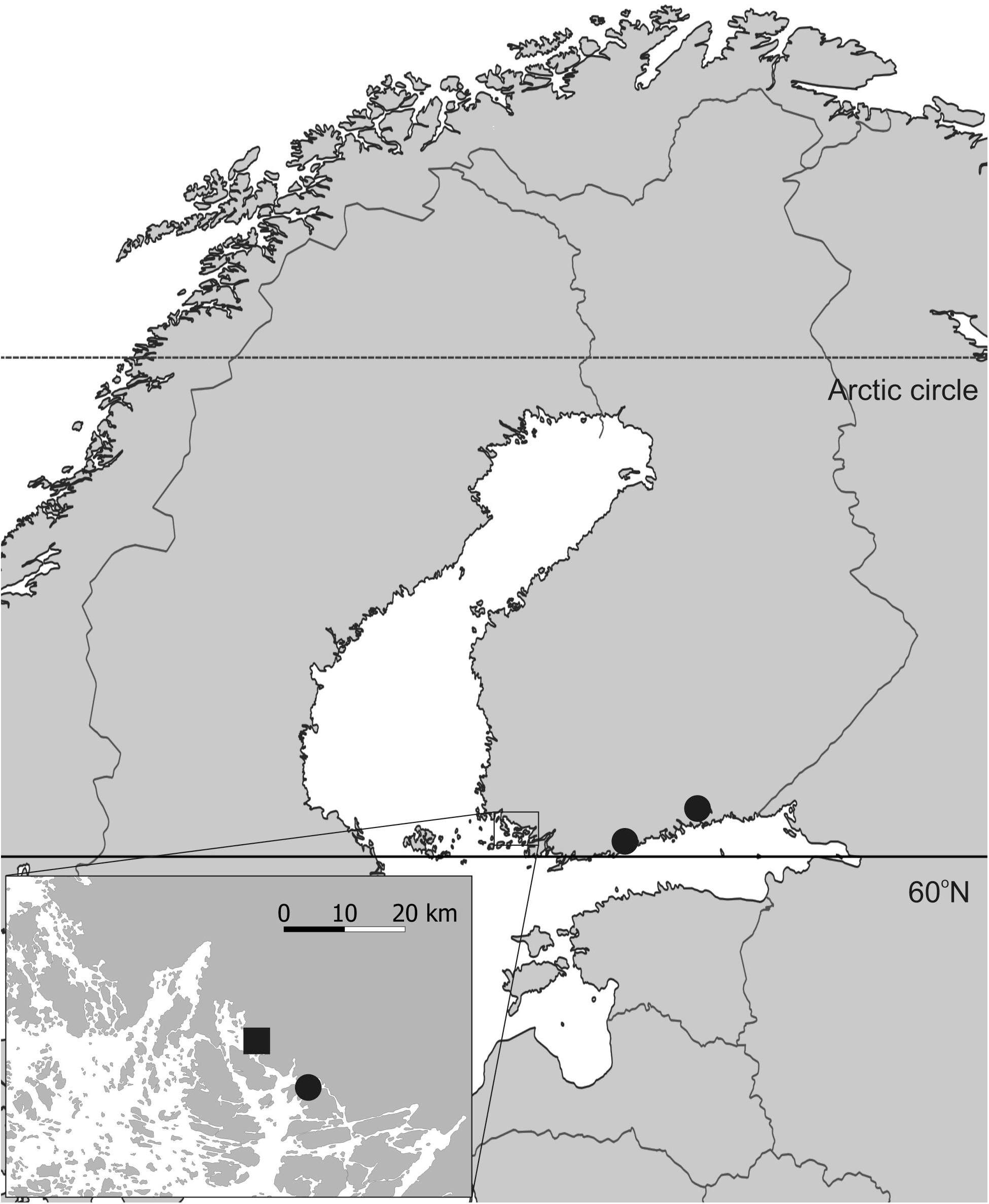
Known breeding colonies (round symbols) and overwintering site (square symbol) of *P. nathusii* in Finland.

We recorded the echolocation calls of bats at 16 sites in South-Western Finland over two consecutive winters (2017-2018 and 2018-2019) by using SongMeter SM2 + Bat passive ultrasound detectors and SMX-US ultrasound microphones (Wildlife Acoustics). These sites included four outcrop formations, three ancient shores, three glacial erratic or boulder formations, three cellars and three control sites in diverse environments where we did not expect bats to be hibernating. We filtered the data using Kaleidoscope Pro (Wildlife Acoustics) and identified the *P. nathusii* calls manually. In our data, the echolocation calls of *P. nathusii* occurred regularly at a single site, Härmälä cleft, over the winter of 2018–2019, suggesting that the species overwintered at the site. Härmälä cleft (N60.488358, E22.006638) is an outcrop formation on the shore of the Baltic Sea in the Masku municipality, South-Western Finland. The cleft is approximately 20 meters high with multiple deep crevices piercing in to the rock. While winter activity, as evidenced by acoustic data, suggests bats hibernate in the cleft, we have never observed any bats in the cleft or visible in the crevices. At the other 15 sites, no observations of the species were made later than the 5^th^ of November.

Our data revealed echolocation calls of *P. nathusii* at Härmälä cleft on several occasions over the winter of 2018–2019 (Table 1; Figure 2). In addition to being the coldest month of that winter with a mean temperature of –4.4 C°, January was also the only month without any recordings of the species. Our results provide further evidence on the hibernation range expansion of *P. nathusii*. In the summer of 2017, we captured two *P. nathusii* weanlings in Turku, approximately 10 km from Härmälä cleft, indicating that the species is also breeding in the area. Our observations in Finland are consistent with other reports on the exceptional breeding and overwintering range expansion of *P. nathusii*. Furthermore, our finding indicates that acoustic surveillance of hibernation sites can yield valuable information on more elusive bat species. Our results suggest that *P. nathusii* has joined the seven species of bats found to hibernate above 60°N in Fennoscandia.

**Table 1.** Number of minutes with recordings of *P. nathusii* at Härmälä cleft during the winter of 2018–2019.

| **Date** | **Minutes with *P. nathusii* observations** |
| --- | --- |
| **6.10.2018** | 29 |
| **8.10.2018** | 9 |
| **9.10.2018** | 21 |
| **10.10.2018** | 6 |
| **11.10.2018** | 47 |
| **12.10.2018** | 21 |
| **13.10.2018** | 7 |
| **15.10.2018** | 6 |
| **17.10.2018** | 1 |
| **3.11.2018** | 5 |
| **13.11.2018** | 10 |
| **20.12.2018** | 2 |
| **15.2.2019** | 5 |
| **23.3.2019** | 2 |
| **16.4.2019** | 2 |
| **17.4.2019** | 1 |
| **24.4.2019** | 4 |

**Figure 2.**
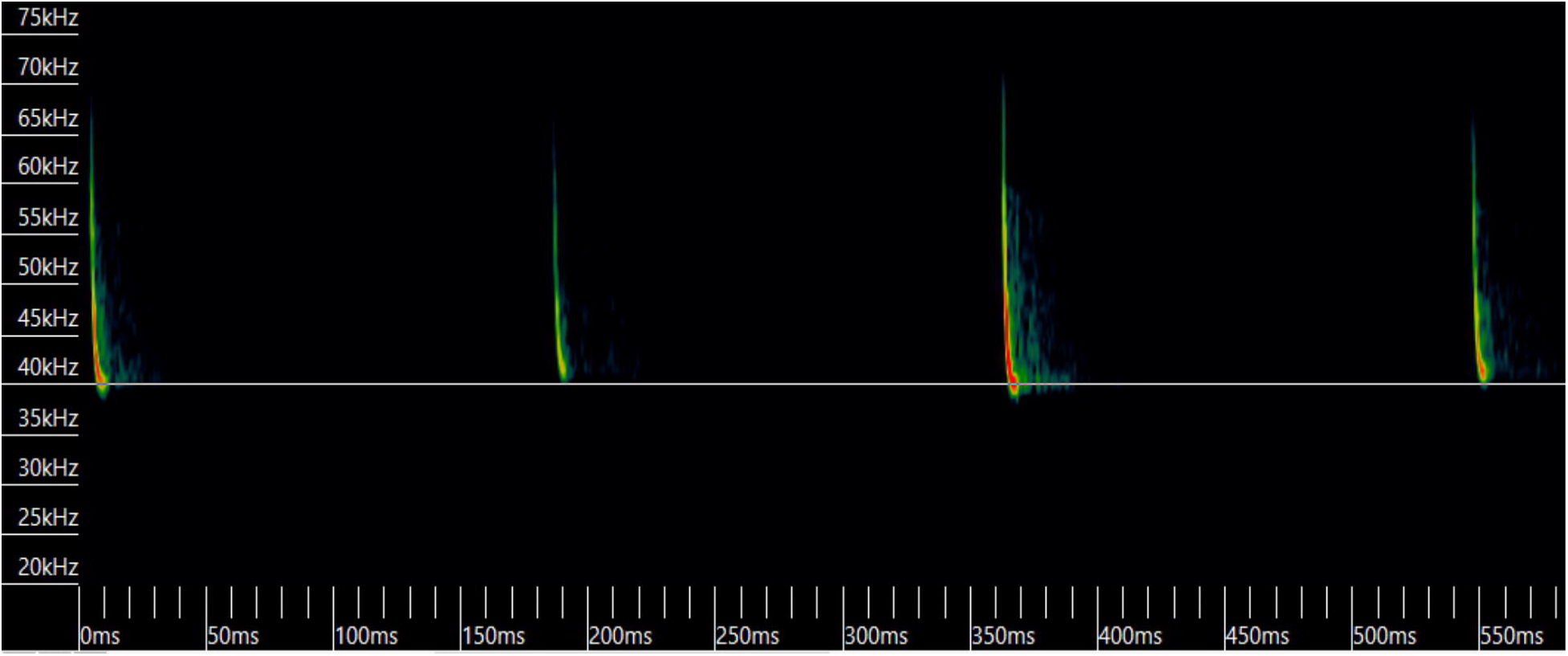
A sonogram of *P. nathusii* calls recorded 15.2.2019.

## Acknowledgements

The study was funded by Kone foundation and H2020 Marie Sklodowska-Curie Actions.

## References

Ancillotto, L., L. Santini, N. Ranc, L. Maiorano and D. Russo. 2016. Extraordinary range expansion in a common bat: the potential roles of climate change and urbanisation. Sci. Nat. 103: 15.

Benda, P. and J. Hotový. 2004. Nález zimujícího netopýra parkového (Pipistrellus nathusii) na jižní Moravě. Vespertilio 8: 137–139. (in Czech)

Brommer, J. E., A. Lehikoinen and J. Valkama. 2012. The Breeding Ranges of Central European and Arctic Bird Species Move Poleward. PLoS ONE 7: e43648.

Diez, C. and A. Kiefer. 2016. Bats of Britain and Europe. Bloomsbury, London, UK.

Flaquer, C., R. Ruiz-Jarillo, I. Torre and A. Arrizabalaga. 2005. First resident population of Pipistrellus nathusii (Keyserling and Blasius, 1839) in the Iberian Peninsula. Acta Chiropterologica 7: 183–188.

Haarsma, A.-J., P. H. C. Lina, A. M. Voûte and H. Siepel. 2019. Male long-distance migrant turned sedentary; The West European pond bat (Myotis dasycneme) alters their migration and hibernation behaviour. PLOS ONE 14: e0217810.

Hagner-Wahlsten, N. and R. Karlsson. 2009. Seurasaaren lepakkoselvitys 2009. Bat house. Helsingin kaupunki, rakennusvirasto. (in Finnish)

Hickling, R., D. B. Roy, J. K. Hill, R. Fox and C. D. Thomas. 2006. The distributions of a wide range of taxonomic groups are expanding polewards. Glob. Change Biol. 12: 450–455.

Humphries, M. M., D. W. Thomas and J. R. Speakman. 2002. Climate-mediated energetic constraints on the distribution of hibernating mammals. Nature 418: 313–316.

Ijäs, A., A. Kahilainen, V. V. Vasko and T. M. Lilley. 2017. Evidence of the Migratory Bat, Pipistrellus nathusii, Aggregating to the Coastlines in the Northern Baltic Sea. Acta Chiropterologica 19: 127–139.

Lane, J. E., L. E. B. Kruuk, A. Charmantier, J. O. Murie and F. S. Dobson. 2012. Delayed phenology and reduced fitness associated with climate change in a wild hibernator. Nature 489: 554–557.

Lehotská, B. and R. Lehotský. 2006. First record of Hypsugo savii (Chiroptera) in Slovakia. Biologia (Bratisl.) 61: 192.

Levinsky, I., F. Skov, J.-C. Svenning and C. Rahbek. 2007. Potential impacts of climate change on the distributions and diversity patterns of European mammals. Biodivers. Conserv. 16: 3803–3816.

Lundy, M., I. Montgomery and J. Russ. 2010. Climate change-linked range expansion of Nathusius’ pipistrelle bat, Pipistrellus nathusii (Keyserling & Blasius, 1839). J. Biogeogr. 37: 2232–2242.

Martinoli, A., D. G. Preatoni and G. Tosi. 2000. Does Nathusius’ pipistrelle Pipistrellus nathusii (Keyserling & Blasius, 1839) breed in northern Italy? J. Zool. 250: 217–220.

Matthews, F., L. M. Kubasiewicz, J. Gurnell, C. A. Harrower, R. A. McDonald and R. F. Shore. 2018. Review of the Population and Conservation Status of British Mammals. A report by the Mammal Society under contract to Natural England, Natural Resources Wales and Scottish Natural Heritage. Natural England, Peterborough.

Mikkonen, S., M. Laine, H. M. Mäkelä, H. Gregow, H. Tuomenvirta, M. Lahtinen and A. Laaksonen. 2015. Trends in the average temperature in Finland, 1847-2013. Stoch. Environ. Res. Risk Assess. 29: 1521–1529.

Newson, S., S. Mendes, H. Crick, N. Dulvy, J. Houghton, G. Hays, A. Hutson, C. Macleod, G. Pierce and R. Robinson. 2009. Indicators of the impact of climate change on migratory species. Endanger. Species Res. 7: 101–113.

Nusová, G., M. Uhrin and P. Kaňuch. 2019. Go to the city: urban invasions of four pipistrelle bat species in eastern Slovakia. Eur. J. Ecol. 5: 23–26.

Parmesan, C. and G. Yohe. 2003. A globally coherent fingerprint of climate change impacts across natural systems. Nature 421: 37–42.

Pētersons, G. 2004. Seasonal migrations of north-eastern populations of Nathusius bat Pipistrellus nathusii (Chiroptera). Myotis 41–42: 29–56.

Rebelo, H., P. Tarroso and G. Jones. 2010. Predicted impact of climate change on European bats in relation to their biogeographic patterns. Glob. Change Biol. 16: 561–576.

Russ, J. M., A. M. Hutson, W. I. Montgomery, P. A. Racey and J. R. Speakman. 2001. The status of Nathusius’ pipistrelle (Pipistrellus nathusii Keyserling & Blasius, 1839) in the British Isles. J. Zool. 254: 91–100.

Rydell, J., L. Bach, P. Bach, L. G. Diaz, J. Furmankiewicz, N. Hagner-Wahlsten, E.-M. Kyheröinen, T. Lilley, M. Masing, M. M. Meyer, G. Pētersons, J Šuba, V. Vasko, V. Vintulis and A. Hedenström. 2014. Phenology of migratory bat activity across the Baltic Sea and the south-eastern North Sea. Acta Chiropterologica 16: 139–147.

Sachanowicz, K. and M. Ciechanowski. 2006. First winter record of the migratory bat Pipistrellus nathusii (Keyserling and Blasius 1839) (Chiroptera: Vespertilionidae) in Poland: yet more evidence of global warming? Mammalia 70: 168–169.

Sachanowicz, K., M. Ciechanowski, P. Tryjanowski and J. Z. Kosicki. 2019. Wintering range of Pipistrellus nathusii (Chiroptera) in Central Europe: has the species extended to the north-east using urban heat islands? Mammalia 83: 260–271.

Sachanowicz, K., A. Wower and A.-T. Bashta. 2006. Further range extension of Pipistrellus kuhlii (Kuhl, 1817) in Central and Eastern Europe. Acta Chiropterologica 8: 543–548.

Sherwin, H. A., W. I. Montgomery and M. G. Lundy. 2013. The impact and implications of climate change for bats: Bats and climate change. Mammal Rev. 43: 171–182.

Thomas, C. D. 2010. Climate, climate change and range boundaries: Climate and range boundaries. Divers. Distrib. 16: 488–495.

Thomas, C., A. Franco and J. Hill. 2006. Range retractions and extinction in the face of climate warming. Trends Ecol. Evol. 21: 415–416.

Tingley, M. W., W. B. Monahan, S. R. Beissinger and C. Moritz. 2009. Birds track their Grinnellian niche through a century of climate change. Proc. Natl. Acad. Sci. 106: 19637–19643.

Uhrin, M., U. Hüttmeir, M. Kipson, P. Estók, K. Sachanowicz, S. Bücs, B. Karapandža, M. Paunović, P. Presetnik, A.-T. Bashta, E. Maxinová, B. Lehotská, R. Lehotský, L. Barti, I. Csösz, F. Szodoray-Paradi, I. Dombi, T. Görföl, S. A. Boldogh, C. Jére, I. Pocora and P. Benda. 2016. Status of Savi’s pipistrelle H ypsugo savii (Chiroptera) and range expansion in Central and South-Eastern Europe: a review: Range changes in Savi’s pipistrelle. Mammal Rev. 46: 1–16.

Warren, M. S., J. K. Hill, J. A. Thomas, J. Asher, R. Fox, B. Huntley, D. B. Roy, S. G. Willis and J. N. Greatorex-Davies. 2001. Rapid responses of British butterflies to opposing forces of climate and habitat change. Nature 414: 65–68.

Willis, C. K. R. and R. M. Brigham. 2003. New Records of the Eastern Red Bat, Lasiurus borealis, from Cypress Hills Provincial Park, Saskatchewan: A Response to Climate Change? Can. Field-Nat. 117: 651–654.

